# Controlled near-space exposure induces community restructuring in an ocean water microbiome

**DOI:** 10.64898/2026.09.11.750916

**Authors:** Huan Gu, Chong Qiu, Tarek S Ibrahim

## Abstract

Marine microbiomes may experience extreme environmental stress during atmospheric transport, yet how near-space conditions reshape natural microbial communities remains poorly understood. Here, we investigated the effects of controlled near-space exposure on a seawater microbiome using high-altitude balloon experiments and 16S rRNA amplicon sequencing. While alpha diversity remained relatively stable across exposure conditions, weighted UniFrac and PERMANOVA analyses revealed reproducible shifts in microbial community composition associated with flight exposure. Taxonomic profiling demonstrated coordinated restructuring across multiple microbial lineages rather than dominance of a single taxon-specific response. Comparable beta-dispersion among groups supported the robustness of these compositional differences. Together, these findings demonstrate that near-space exposure was associated with reproducible restructuring of microbial community composition without substantial loss of overall diversity. Raw sequencing data have been deposited in the NCBI Sequence Read Archive under BioProject accession PRJNA1479017.

## 1. Introduction

Marine microbiomes are major drivers of global biogeochemical cycling and ecosystem function in the surface ocean(*1–4*). These communities are highly sensitive to environmental stressors, including ultraviolet (UV) radiation, low temperature, oxidative stress, and desiccation(*5–7*). Because microbial taxa differ substantially in stress tolerance and DNA repair capacity, extreme environmental gradients may act as deterministic ecological filters that reshape microbial community assembly and diversity(*8*).

Marine microorganisms can enter the atmosphere through sea spray aerosolization driven by bubble bursting and ocean-atmosphere exchange processes(*9–12*). Once aerosolized, microbes may experience severe near-space stressors including elevated UV radiation, low pressure, and extreme cold during transport into the upper troposphere and lower stratosphere(*13–16*). Previous atmospheric sampling and laboratory simulation studies have demonstrated diverse microbial responses to these conditions(*13, 17–19*). However, how natural marine microbiomes respond at the community level to authentic near-space exposure remains poorly understood.

High-altitude balloon platforms provide a practical approach for exposing biological samples to real near-space environmental conditions(*15, 20*). While previous studies have primarily focused on isolated microorganisms or spores, the ecological responses of complex marine microbiomes remain largely unexplored. In particular, whether near-space conditions impose selective ecological filtering capable of restructuring natural marine microbial communities has not been comprehensively investigated. Marine microorganisms embedded within micron-scale cloud droplets may additionally experience partial protection from desiccation, potentially altering microbial responses during atmospheric transport(*11, 21, 22*).

Here, we investigated the effects of controlled short-duration near-space exposure on a natural seawater microbiome using 16S rRNA amplicon sequencing. Our experimental design incorporated matched ground controls together with shielded and unshielded flight samples to distinguish exposure-driven ecological filtering from handling effects. We hypothesized that near-space exposure would selectively restructure marine microbiome diversity and composition, and that shielding from direct environmental exposure would partially attenuate these effects. By focusing on community-level assembly responses rather than survival of individual taxa, this study provides experimental evidence that extreme atmospheric gradients can selectively reshape marine microbial ecosystems.

## 2. Results

### 2.1 Robust sequencing and feature curation support reliable ecological comparisons

To evaluate microbiome responses across Flight_UV, Flight_UVshield, Ground_pre, and Ground_post conditions, we implemented a standardized 16S rRNA amplicon sequencing workflow incorporating denoising, chimera removal, contamination assessment, and prevalence-based feature curation (Fig. 1). Sequencing quality-control analyses demonstrated consistent sequencing depth and stable read retention across samples (Supplementary Figs. S1–S2). Negative-control analyses revealed substantially lower taxonomic complexity and distinct abundance patterns relative to biological samples, indicating limited contribution of background contamination to the dominant ecological structure of the dataset (Supplementary Fig. S3). Furthermore, prevalence-based filtering substantially reduced low-abundance features while preserving the major ecological structure of the dataset (Supplementary Fig. S4). Together, these results support the robustness of downstream diversity, compositional, and differential abundance analyses and indicate that observed community differences primarily reflect biological responses to near-space exposure rather than technical artifacts.

**Figure 1.**
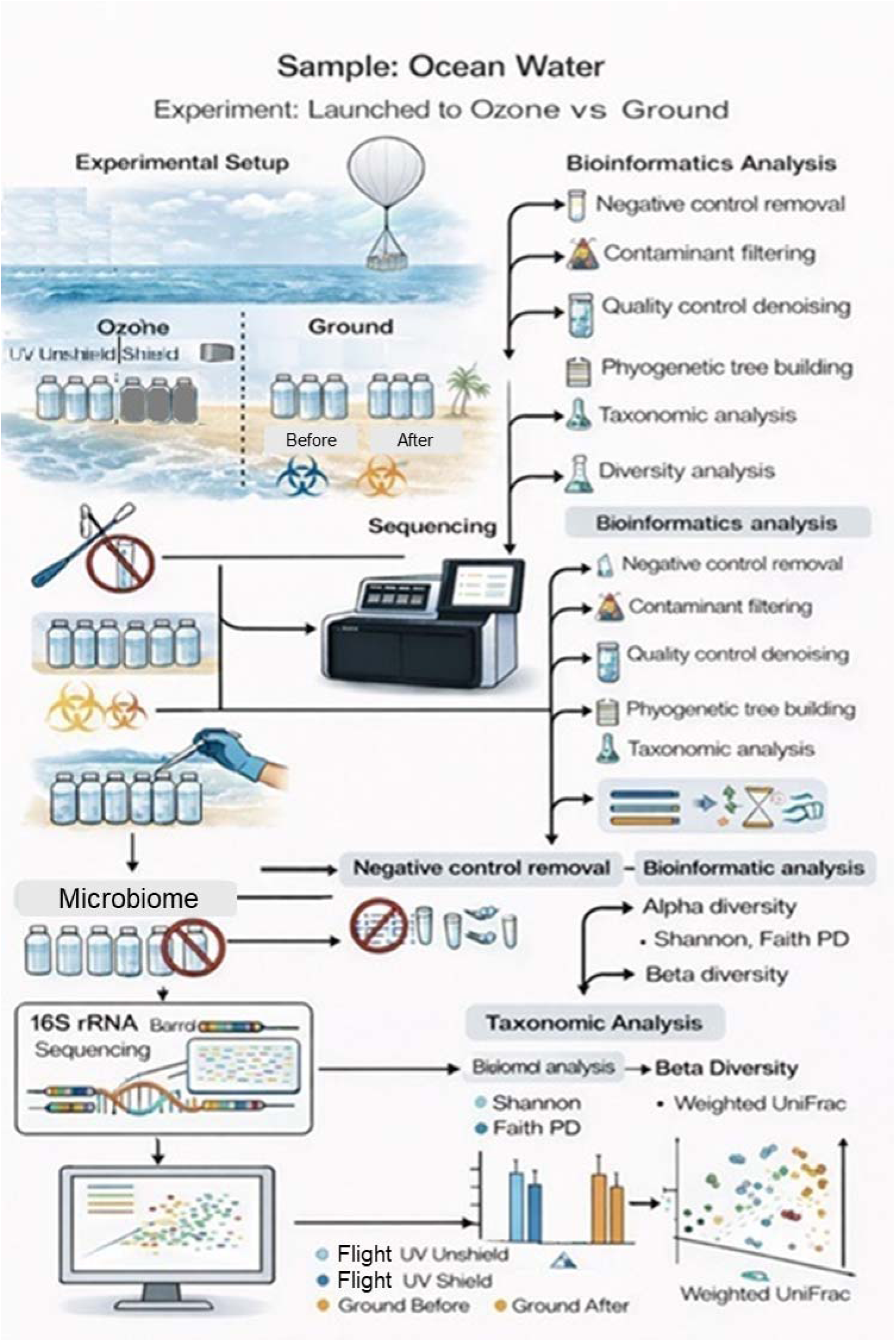
Experimental workflow and analytical overview. Schematic representation of the sequencing and bioinformatic pipeline used to characterize microbial community responses across near-space exposure conditions. Raw reads underwent denoising, chimera removal, and prevalence-based feature filtering prior to diversity analysis, multivariate statistical testing, and taxonomic profiling. Supplementary figures provide detailed quality-control validation for each analytical step.

### 2.2 Near-space exposure preserves overall microbial diversity

We first evaluated whether near-space exposure altered overall microbial diversity. Shannon diversity and Faith’s phylogenetic diversity exhibited limited variation across Flight_UV, Flight_UVshield, Ground_pre, and Ground_post conditions following rarefaction normalization (Fig. 2A–B), indicating that overall richness and phylogenetic breadth were largely preserved. Rarefaction analyses confirmed sufficient sequencing depth and stable diversity estimation across samples (Supplementary Fig. S2). Together, these results suggest that near-space exposure did not induce broad-scale diversity loss, but instead may have driven more selective compositional restructuring within the microbial community.

**Figure 2.**
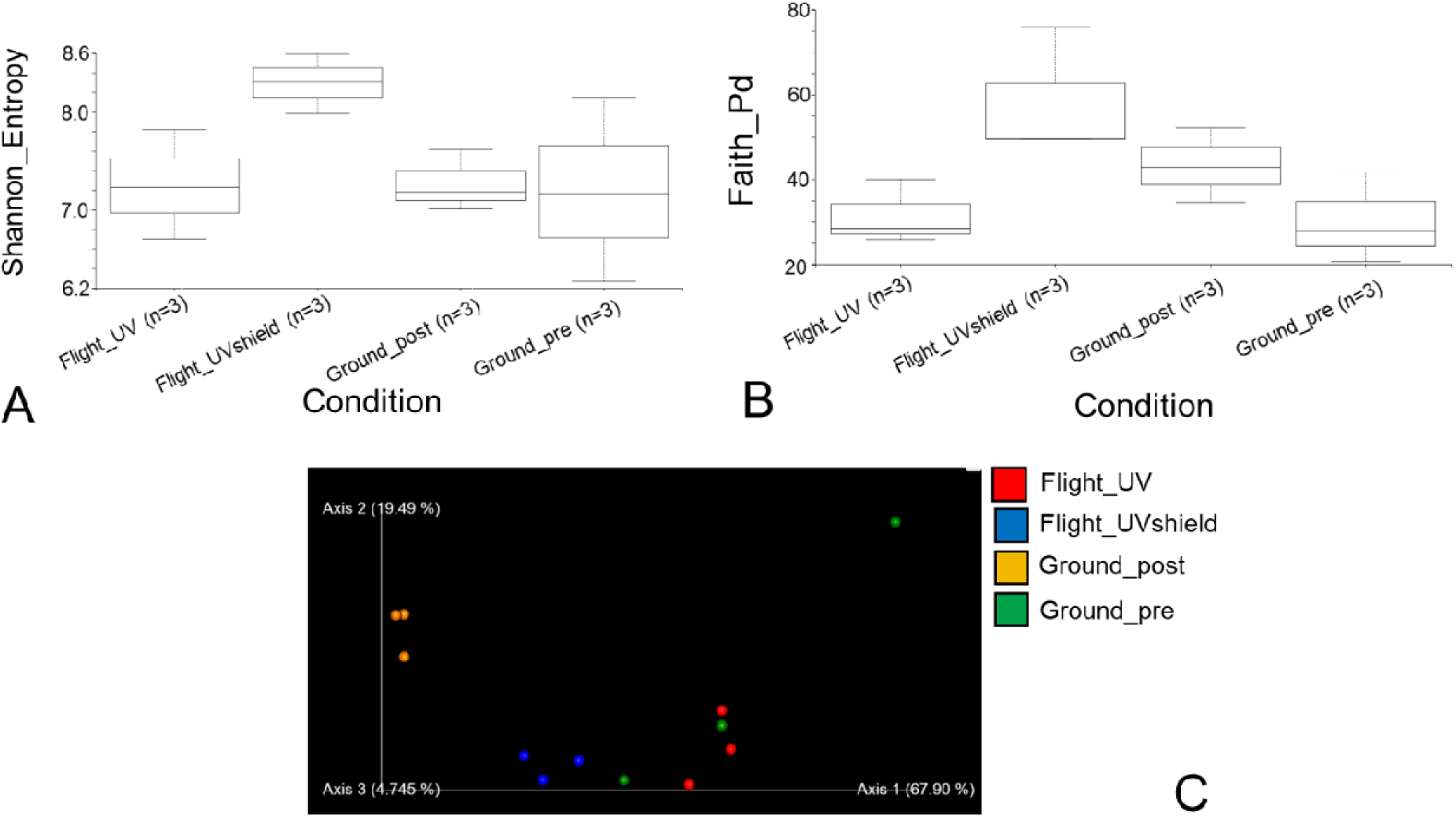
Diversity metrics and phylogenetic ordination of microbial communities following near-space exposure. (A) Shannon diversity and (B) Faith’s phylogenetic diversity across experimental conditions following rarefaction normalization. (C) Weighted UniFrac principal coordinates analysis (PCoA) showing condition-dependent restructuring of microbial community composition across flight-exposed and ground-control samples.

### 2.3 Phylogenetic ordination reveals reproducible restructuring of microbial community composition

Although alpha diversity remained relatively stable, beta-diversity analysis revealed clear shifts in microbial community organization across exposure conditions. Principal coordinates analysis (PCoA) based on weighted UniFrac distances demonstrated separation between flight-exposed samples and ground controls along the major axes of variation (Fig. 2C), consistent with phylogenetically structured community restructuring associated with near-space exposure. Replicate samples clustered reproducibly within experimental groups, supporting the robustness of these compositional differences

### 2.4 Alpha diversity

Alpha-diversity analyses revealed that near-space exposure produced relatively modest effects on within-sample microbial diversity. Shannon diversity and Faith’s phylogenetic diversity varied among experimental conditions following rarefaction normalization (Fig. 2A–B), indicating that near-space exposure influenced the relative distribution and phylogenetic composition of microbial taxa. Rarefaction analyses demonstrated that diversity estimates approached saturation before the selected sequencing depth threshold, supporting the robustness of downstream diversity comparisons (Supplementary Fig. S2)

Despite these differences, overall richness and phylogenetic breadth were largely preserved across Flight_UV, Flight_UVshield, Ground_pre, and Ground_post communities. Collectively, these findings suggest that near-space exposure did not induce broad-scale diversity loss, but instead was associated with more subtle changes in community structure that were better reflected by community composition analyses than by richness-based diversity metrics

### 2.5 Taxonomic profiling reveals coordinated restructuring across multiple microbial lineages

Although alpha diversity remained relatively stable, beta-diversity analyses revealed clear shifts in microbial community organization across exposure conditions. Principal coordinates analysis (PCoA) based on weighted UniFrac distances demonstrated separation between flight-exposed samples and ground controls along the major axes of variation (Fig. 2C), indicating that near-space environmental conditions altered the phylogenetic structure of the microbial community.

Replicate samples clustered reproducibly within experimental groups, supporting the consistency of the observed compositional patterns. Both Flight_UV and Flight_UVshield communities diverged from Ground_pre and Ground_post controls, suggesting that community restructuring was associated with the combined effects of near-space environmental stressors rather than ultraviolet radiation alone. Together, these results indicate that near-space exposure influenced microbial community assembly through compositional reorganization despite relatively limited changes in overall diversity.

### 2.6 Multivariate statistical validation confirms exposure-associated community restructuring

To statistically evaluate the compositional shifts observed in ordination space, permutational multivariate analysis of variance (PERMANOVA) was performed using weighted UniFrac distance matrices. Global analysis identified significant differences in microbial community composition among experimental conditions (PERMANOVA, p = 0.001; 999 permutations), supporting the separation patterns observed in the weighted UniFrac ordination (Fig. 3A).

**Figure 3.**
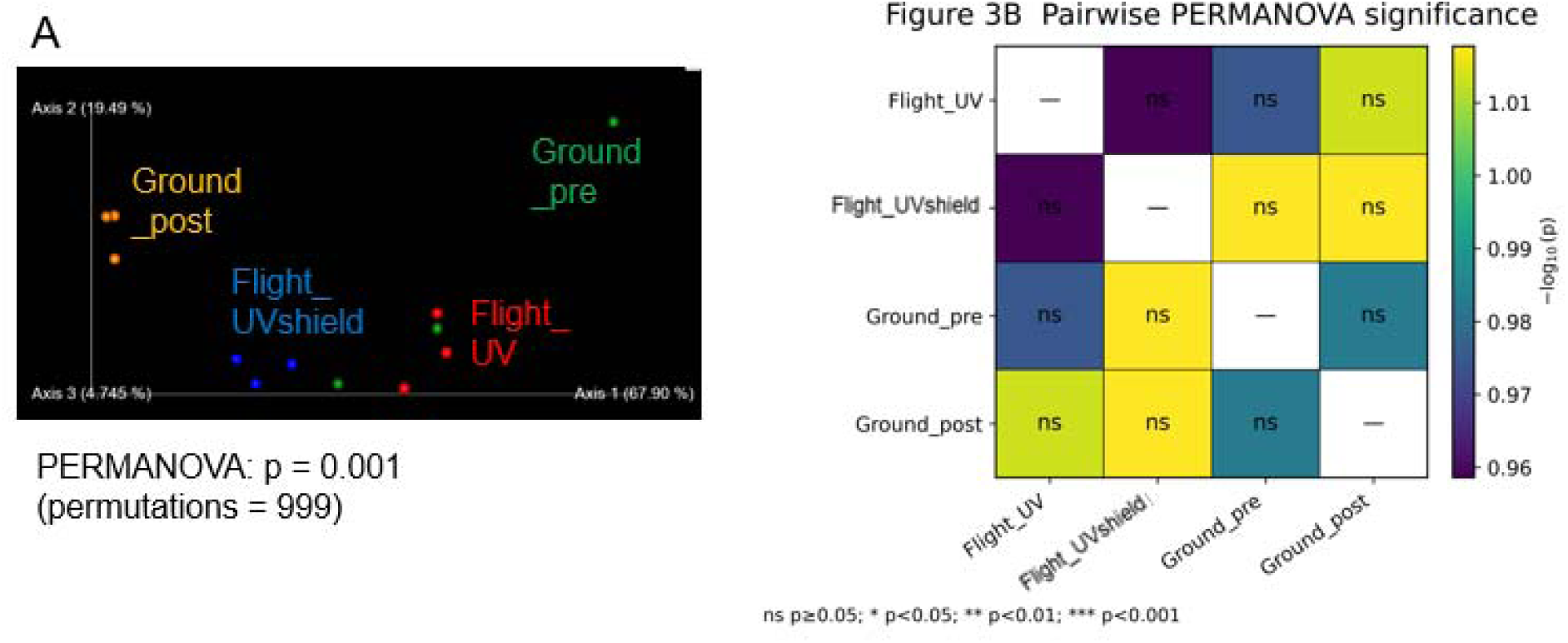
Statistical validation of microbial community restructuring following near-space exposure. (A) Principal coordinates analysis annotated with PERMANOVA results demonstrating significant differences in microbial community composition across experimental conditions (p = 0.001; 999 permutations). (B) Heatmap summarizing pairwise PERMANOVA comparisons among experimental groups. Cell color represents −log10(p) values, and annotations indicate permutation-derived significance levels (ns, p ≥ 0.05; *p < 0.05; **p < 0.01; ***p < 0.001). The predominance of non-significant pairwise contrasts suggests distributed multigroup compositional restructuring rather than isolated pairwise divergence.

Pairwise PERMANOVA comparisons revealed variable degrees of divergence among flight and ground conditions, although most individual contrasts did not reach statistical significance (Fig. 3B; Table S1). This pattern suggests that near-space exposure induced distributed multigroup restructuring of microbial communities rather than a dominant shift between any single pair of conditions. Importantly, beta-dispersion analyses demonstrated comparable within-group dispersion across experimental conditions (Supplementary Fig. S5), indicating that the observed compositional differences were not driven by heterogeneous variance among groups. Collectively, these findings provide statistical support for exposure-associated restructuring of microbial community composition.

### 2.7 Taxonomic profiling reveals coordinated restructuring across microbial lineages

To identify the taxonomic basis underlying the compositional restructuring observed in diversity analyses, microbial community composition was examined across hierarchical taxonomic levels. Dominant bacterial phyla were generally conserved across experimental conditions, although relative abundances of multiple taxa varied between flight-exposed and ground-control samples (Fig. 4A).

**Figure 4.**
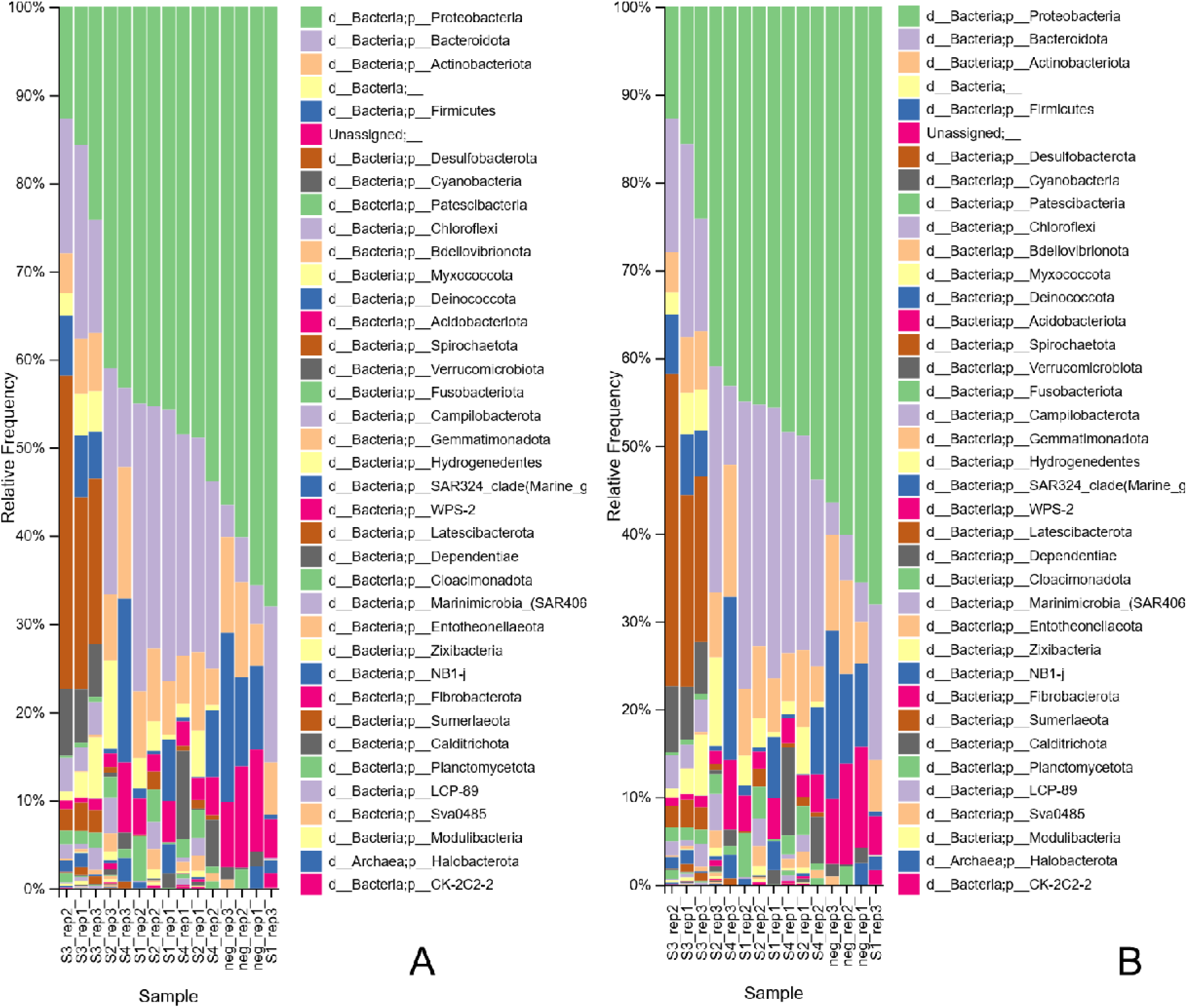
Taxonomic composition of microbial communities across experimental conditions. (A) Relative abundance of dominant bacterial phyla across samples. (B) Genus-level composition showing distributed compositional restructuring across multiple microbial lineages associated with near-space exposure. S1: Flight_UV; S2: Flight_UVShield; S3:Ground_post; S4:Ground_pre; neg: No flight and UV.

At the genus level, compositional differences were distributed across diverse microbial lineages rather than being driven by expansion or depletion of a single dominant taxon (Fig. 4B). Taxonomic assignment depth varied across features, with a substantial proportion of sequences classified to the genus level and a smaller fraction remaining unresolved (Supplementary Fig. S5). These observations indicate that near-space exposure was associated with coordinated restructuring across multiple microbial groups rather than isolated changes within individual taxa.

### 2.8 Differential abundance analysis identifies taxa associated with near-space exposure

Differential abundance analysis using ANCOM identified multiple taxa exhibiting strong compositional shifts across experimental conditions (Fig. 5; Table S2). Several genera, including Hyphomonas, Sphingomicrobium, Desulfoconvexum, Desulforhopalus, Nocardioides, and Rubrivirga, displayed among the highest ANCOM W statistics, indicating consistent abundance differences across pairwise log-ratio comparisons. The twenty taxa with the highest W statistics are summarized in Supplementary Table S2.

**Figure 5.**
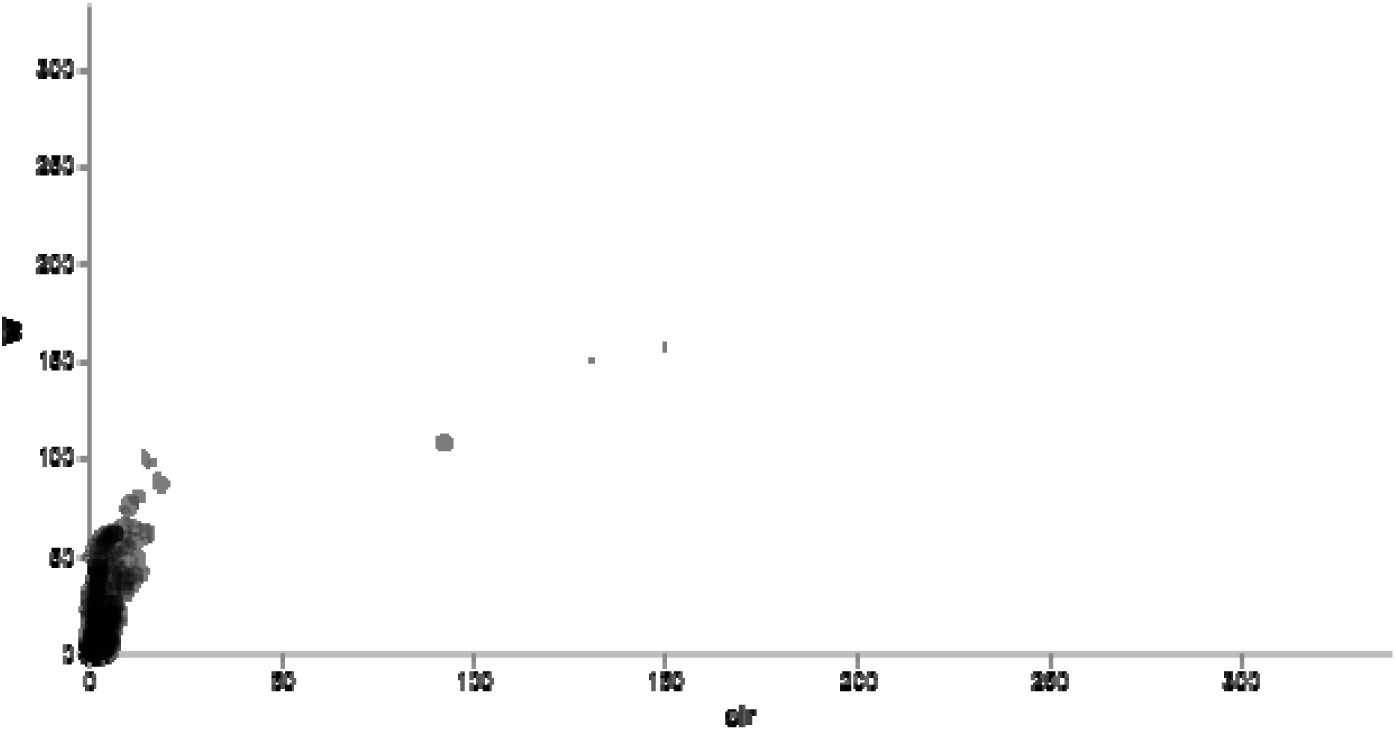
Differential abundance analysis of microbial taxa across experimental conditions using ANCOM. Each point represents a taxon. The W statistic reflects the number of pairwise log-ratio tests in which the null hypothesis of equal abundance was rejected. Taxa with high W values exhibit the strongest evidence for differential abundance among experimental conditions. Multiple phylogenetically distinct taxa displayed elevated W statistics, consistent with distributed community restructuring following near-space exposure.

Notably, the taxa identified by ANCOM spanned multiple phylogenetic lineages rather than being restricted to a single taxonomic group. This pattern suggests that near-space exposure influenced microbial community composition through distributed ecological restructuring rather than selective enrichment or depletion of a single dominant taxon. Consistent with beta-diversity and PERMANOVA analyses, differential abundance signals were broadly distributed across the community, supporting the hypothesis that near-space exposure acted as an ecological filter that altered the relative representation of multiple microbial lineages. Collectively, these findings indicate that community-level restructuring was driven by coordinated compositional shifts across diverse taxa rather than by changes in a limited subset of dominant organisms.

## 3. Discussion

Environmental microbiomes often respond to physical stress through gradual community restructuring rather than abrupt compositional collapse. In this study, near-space exposure was associated with reproducible shifts in microbial community composition despite relatively stable alpha diversity, consistent with previous observations that environmental constraints can alter community composition without necessarily reducing overall diversity(*8, 14, 16, 23*). Ordination and PERMANOVA analyses demonstrated exposure-associated community divergence (Fig. 2C; Fig. 3A), while the predominance of non-significant pairwise contrasts (Fig. 3B; Table S1) suggests that the observed differences were distributed across multiple experimental groups rather than driven by a single dominant transition. Furthermore, comparable within-group dispersion across experimental conditions (Supplementary Fig. S6) supports the interpretation that these differences reflect compositional restructuring rather than variance-driven statistical artifacts(*24, 25*).

The moderate but consistent separation observed across exposure conditions is consistent with previous studies showing that radiation exposure, confinement, and altered environmental conditions can reshape microbial community composition under extreme environmental conditions(*13, 14, 16*). Notably, both UV-exposed and shielded flight samples exhibited compositional divergence from ground controls (Fig. 2C; Fig. 3A), suggesting that microbial responses likely reflected the combined influence of multiple near-space environmental factors rather than a single isolated stressor. Such distributed community responses are increasingly recognized in environmental microbiome systems, where subtle environmental changes can produce measurable beta-diversity shifts without major taxonomic collapse(*11, 13*).

Taxonomic profiling and differential abundance analyses further supported this interpretation. Community differences were distributed across multiple microbial lineages rather than driven by expansion or depletion of a single dominant taxon (Fig. 4A–B). ANCOM analysis identified multiple genera exhibiting elevated W statistics, including Hyphomonas, Sphingomicrobium, Desulfoconvexum, Desulforhopalus, Nocardioides, and Rubrivirga (Fig. 5; Table S2), providing feature-level evidence for exposure-associated differences in relative abundance. Importantly, these taxa span diverse phylogenetic groups, indicating that the observed responses were broadly distributed throughout the community rather than concentrated within a single lineage. Variable assignment depth at lower taxonomic ranks (Supplementary Fig. S5) additionally reflects the inherent classification limitations associated with complex environmental microbiomes and incomplete reference databases(*17, 18*).

Because low-biomass microbiome studies are particularly susceptible to background contamination, contamination control was incorporated throughout sample processing and bioinformatic analysis. Extraction and handling controls were processed alongside biological samples, and contaminant assessment was performed using abundance and prevalence patterns rather than taxonomy alone. Negative controls exhibited substantially lower taxonomic complexity than biological samples (Supplementary Fig. S3), while prevalence-based filtering removed low-frequency features without altering the major ecological structure of the dataset (Supplementary Fig. S4). Furthermore, the compositional differences identified through ordination, PERMANOVA, and ANCOM analyses remained evident following contaminant filtering, supporting the interpretation that the observed community patterns primarily reflect biological variation associated with exposure conditions rather than laboratory-derived background signal.

Together, these findings demonstrate that near-space exposure was associated with reproducible restructuring of marine microbial community composition while preserving overall diversity. This pattern is consistent with broader environmental microbiome studies showing that physical stressors can alter microbial community composition without inducing large-scale diversity loss(*13, 14, 16, 23*). Such resilience may reflect functional redundancy and ecological plasticity within marine microbial communities exposed to fluctuating environmental gradients, although the functional consequences of the observed compositional changes remain unresolved.

Several limitations should nevertheless be considered. The present study evaluated compositional responses using 16S rRNA amplicon sequencing and therefore does not directly resolve functional activity or metabolic adaptation. In addition, the seawater samples were designed to simulate micron-scale cloud droplets containing marine microbiomes but did not fully reproduce atmospheric chemical transformations, including exposure to ozone, NOx radicals, or dehydration–condensation cycling. Future studies integrating aerosolized seawater particles, metagenomics, metatranscriptomics, and longitudinal exposure experiments may further clarify the mechanisms underlying near-space-associated community restructuring.

## 4. Conclusions

By integrating phylogenetic ordination, multivariate statistical testing, differential abundance analysis, and dispersion controls, this study provides experimental evidence that near-space exposure is associated with reproducible restructuring of natural marine microbial communities. Despite relatively stable alpha diversity, flight-exposed communities exhibited significant compositional differences from ground controls, indicating that near-space environmental conditions influenced microbial community composition without causing substantial diversity loss. These findings advance understanding of how extreme atmospheric environments affect natural marine microbiomes and highlight the value of combining contamination-aware microbiome workflows with complementary ecological and statistical analyses when evaluating microbial responses to complex environmental stressors. Future studies incorporating larger sample sizes and functional approaches will be important for determining the mechanisms and ecological consequences underlying the compositional changes observed here.

## 5. Materials and Method

### 5.1 Seawater Sampling and Balloon Launch Procedure

Seawater used for microbiome analysis was collected from Long Island Sound (LIS) at Sandy Point Bird Sanctuary (CT) in the evening prior to balloon launch. Using the field measurement data(*26*) for 2024, the pH of surface seawater in LIS during our sampling period was estimated as (7.9 ± 0.1) and its total alkalinity as (1826 ± 31) µmol/kg [corresponding to a salinity of (25.5 ± 0.4) g/kg using the model describing the correlation between total alkalinity and salinity of surface water in Atlantic Ocean(*27, 28*)].

All sampling containers and tools were sterilized by ultraviolet irradiation for 1 hour to minimize contamination. On the morning of launch, the seawater container was gently mixed to ensure homogeneity before subsampling. Baseline microbiome samples (Ground Before, n = 3; 50 mL each) were collected prior to launch by filtration through Millipore® Sterivex™ pressure filters (0.45 μm pore size) using sterile syringes and needles, followed by preservation in RNAlater®. For near-space exposure experiments, six 50 mL aliquots were transferred into sterile sealed IV bags to serve as balloon payload samples. Three payloads were exposed to ultraviolet radiation during flight (Flight_UV), while three were wrapped with UV-shielding material (Flight_UVshield) to isolate non-UV environmental stressors. Remaining seawater was preserved under UV-shielded conditions at room temperature (∼20 °C) to serve as matched ground controls collected after recovery (Ground After). The experimental design and exposure conditions are summarized in Table 1. Exposure conditions will be evaluated using biological replicates (n = 3 per group), allowing application of permutation-based multivariate statistics while maintaining a feasible experimental scale for a one-year environmental analog study.

**Table 1.** Experimental Conditions for Seawater Microbiome Exposure.

| Sample | Sealed | UV | Near-Space Condition |
| --- | --- | --- | --- |
| Ground before (n=3) | × | × | × |
| Flight_UV (n=3) | √ | √ | √ |
| Flight_UVshield (n=3) | √ | × | √ |
| Ground after (n=3) | √ | × | × |

Following balloon recovery, microbiomes from flight-exposed and ground-control samples were harvested using identical filtration procedures and preserved in RNA*later*^®^. Negative microbiome controls consisting of RNA*later*® alone, syringes, needles, filters, and empty containers were processed in parallel to monitor potential background contamination introduced during handling. All filters were transferred into sterile tubes and rapidly frozen at −80 °C until DNA extraction and sequencing.

The weather balloon platform and flight information were described in detail elsewhere(*13*). Our scientific payload design consists of sealed IV bags mounted in series along a lightweight string system, along with a multiband tracking subsystem and sensors for pressure and temperature(*26*). This payload design enabled rapid field deployment while minimizing environmental contamination and allowing controlled comparison between ultraviolet-exposed and UV-shielded samples.

On June 5^th^ 2024, a 1600 g weather balloon launched from West Hartford, Connecticut (41.79635°, –72.7161667°) ascended at an average rate of 4.45 m/s to maximum altitude of 28,580 m after 107 minutes of flight. After the natural burst of the balloon at the maximum altitude, the payload descended at an average rate of –19.8 m/s to the landing location in Willington, Connecticut (41.84415°, –72.24405°) for approximately 24 minutes. The total flight time was 131 minutes. After analysis of the temperature profile during the flight, it was estimated that the payload was exposed to a significant freezing condition (defined as temperature ≤ –20 ⁰C) for approximately 90 minutes, or about 2/3 of the total flight time.

### 5.2 Bioinformatic Workflow Overview

A contamination-aware, phylogeny-resolved bioinformatic workflow was implemented to preserve ecological structure while minimizing background signal in low-biomass ocean water microbiome samples exposed to balloon-mediated ozone and ultraviolet (UV) conditions. The pipeline integrated amplicon sequence variant (ASV) inference, taxonomy-guided contamination filtering, phylogenetic reconstruction, rarefaction-based normalization, and multivariate ecological statistics within QIIME 2 (version 2023.9).

Sequence processing and analysis were performed using the QIIME 2 platform (*29*). Amplicon sequence variants were inferred using the DADA2 algorithm, which models sequencing error profiles and removes chimeric reads(*30*). Taxonomic classification was conducted using a naïve Bayes classifier trained on the SILVA 138 reference database (*31*). Representative sequences were aligned with MAFFT and phylogenetic trees were constructed using FastTree (*32, 33*) for downstream phylogenetic diversity analyses. Beta-diversity metrics including weighted and unweighted UniFrac were calculated to evaluate community dissimilarity (*34*). Statistical comparisons of community structure were performed using PERMANOVA (*35*). Contamination filtering followed prevalence-based principles consistent with the decontam framework for low-biomass microbiome studies (*36*), with detailed implementation described in Supplementary Methods S1–S3.

### 5.2 Amplicon Processing and Taxonomic Classification

Raw paired-end amplicon reads were denoised using the DADA2 plugin to infer ASVs, remove chimeric sequences, and correct sequencing errors. Representative sequences were taxonomically annotated using a naïve Bayes classifier trained on the SILVA 138 reference database trimmed to the amplified region. Taxonomic profiles were examined across hierarchical ranks, and genus-level collapsed tables were used for ecological interpretation and contamination evaluation.

### 5.3 Contamination-Aware Filtering

Because high-altitude ocean aerosol samples represent a low-biomass microbiome environment, rigorous contamination control procedures were applied to reduce the influence of reagent- or handling-derived taxa. Negative control samples sequenced alongside biological samples were used as empirical baselines to identify taxa enriched in controls relative to environmental samples. Detailed detection criteria and implementation steps are provided in Supplementary Methods S1–S2. Flagged taxa were removed using taxonomy-based filtering within QIIME 2, and negative control samples were excluded from downstream ecological analyses.

### 5.4 Phylogenetic Reconstruction

Representative ASV sequences were aligned using MAFFT, masked to remove hypervariable positions, and used to construct a phylogenetic tree with FastTree. The resulting tree was midpoint-rooted to support phylogeny-based diversity calculations.

### 5.5 Rarefaction and Diversity Metrics

The filtered feature table was rarefied to 25,000 sequences per sample, corresponding to the minimum sequencing depth retained after contamination filtering. Alpha diversity was evaluated using Shannon diversity and Faith’s phylogenetic diversity. Beta diversity was quantified using weighted UniFrac, unweighted UniFrac, and Bray–Curtis dissimilarity. Principal coordinates analysis (PCoA) was used to visualize community structure across experimental conditions.

### 5.6 Statistical Analysis of Community Structure

Differences in beta diversity among experimental groups (Flight_UV, Flight_UV shield, Ground_pre, Ground_post) were evaluated using PERMANOVA with 999 permutations and pairwise comparisons where applicable. Alpha diversity differences were assessed using non-parametric Kruskal–Wallis tests. Differential abundance analysis was performed using the Analysis of Composition of Microbiomes (ANCOM) method implemented in QIIME 2. Taxa were aggregated at the genus level, a pseudocount was added prior to compositional analysis, and experimental condition was used as the grouping variable. Additional details are provided in Supplementary Methods S7.

## 6. Ackknowledgement.

The project was supported by Nationwide Eclipse Ballooning – Continuation Grant, NSF CAREER (AGS-1847019), National Science Foundation LEAPS-MPS (NSF: 2418289), and University of New Haven Buckman Endowed Fund.

## 7. Data Availability

The 16S rRNA amplicon sequencing data generated during this study have been deposited in the NCBI Sequence Read Archive (SRA) under BioProject accession PRJNA1479017. Raw sequencing reads and associated metadata are publicly available through the NCBI BioProject database.

