## Supplemental Data 1 for "Controlled near-space exposure induces community restructuring in an ocean water microbiome"

1. **Supplementary Methods — Quality Control, Contamination Assessment, and Downstream Ecological Validation**

Supplementary figures are organized to follow the analytical workflow from sequencing quality control through contamination assessment and downstream ecological validation.

1.1 **Supplementary Methods S1. Sequencing Quality Assessment and Read Retention**

To document sequence processing efficiency and confirm comparable data retention across experimental groups, denoising statistics were summarized across all samples. Read counts at each stage of processing (input, filtering, denoising, merging, and non-chimeric reads) were visualized to evaluate potential biases introduced during sequence inference.

Quality-control summaries demonstrate that sequencing depth and retention rates were comparable across Flight_UV, Flight_UV shield, Ground_pre, and Ground_post samples, indicating that downstream diversity patterns were unlikely to arise from uneven sequence loss (Fig. S1).

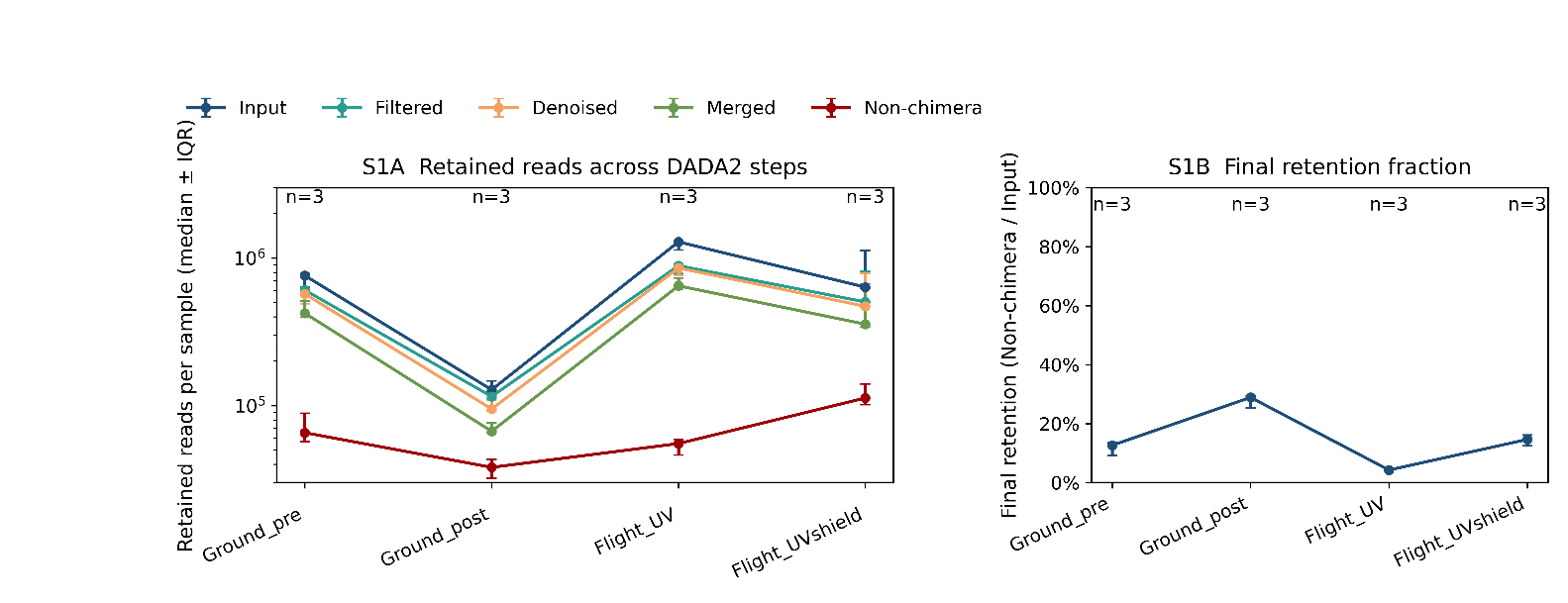

**Figure S1. Read retention during DADA2 processing and final non-chimeric yield.**
(A) Median reads per sample retained across DADA2 steps (Input, Filtered, Denoised, Merged, and Non-chimeric) for each condition. Points indicate medians and error bars show interquartile range (IQR; n = 3 per group; log scale). (B) Final retention fraction, calculated as non-chimeric reads relative to input reads, summarized as median ± IQR (n = 3).

### **1.2 Supplementary Methods S2. Rarefaction Depth Evaluation**

Because rarefaction depth can influence both alpha- and beta-diversity metrics, sequencing depth distributions were examined prior to diversity analysis. Rarefaction curves were generated to evaluate whether diversity estimates approached asymptotic behavior across samples.

Based on these evaluations, a rarefaction depth of 25,000 sequences per sample was selected to balance retention of samples with adequate representation of community diversity while minimizing loss of ecological signal (Fig. S2).

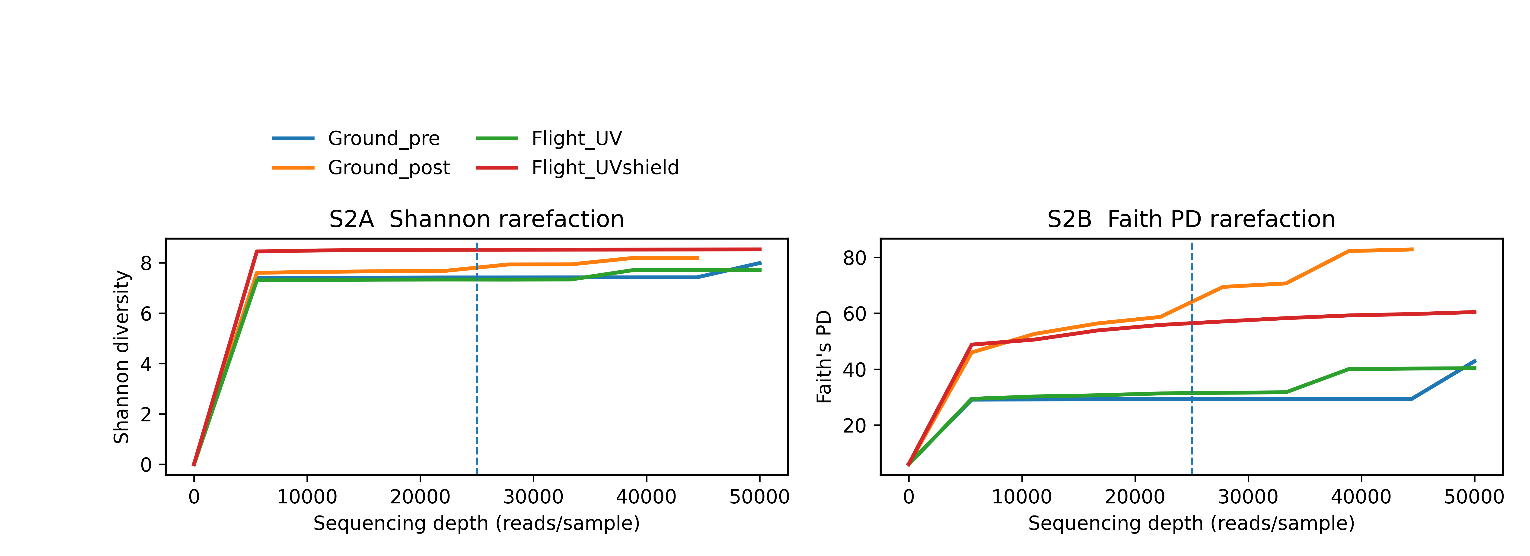

**Figure S2. Rarefaction analysis supporting sequencing depth selection.** (A) Shannon diversity and (B) Faith’s phylogenetic diversity rarefaction curves across sequencing depths, stratified by experimental condition. Curves represent median diversity values across samples following repeated subsampling. Both metrics approach plateau prior to the selected rarefaction depth of 25,000 reads per sample (dashed line), supporting the chosen normalization threshold for downstream diversity analyses.

### **1.3 Supplementary Methods S3. Negative-Control–Guided Contamination Assessment**

Negative controls were included throughout DNA extraction and sequencing to characterize potential background contamination. Rather than removing taxa solely based on taxonomy, contamination evaluation incorporated prevalence and abundance patterns across controls and biological samples.

Visualization of genus-level composition including negative controls allowed identification of taxa enriched in control samples relative to biological ocean water communities. This comparison provided empirical justification for downstream filtering decisions while minimizing removal of legitimate environmental taxa (Fig. S3).

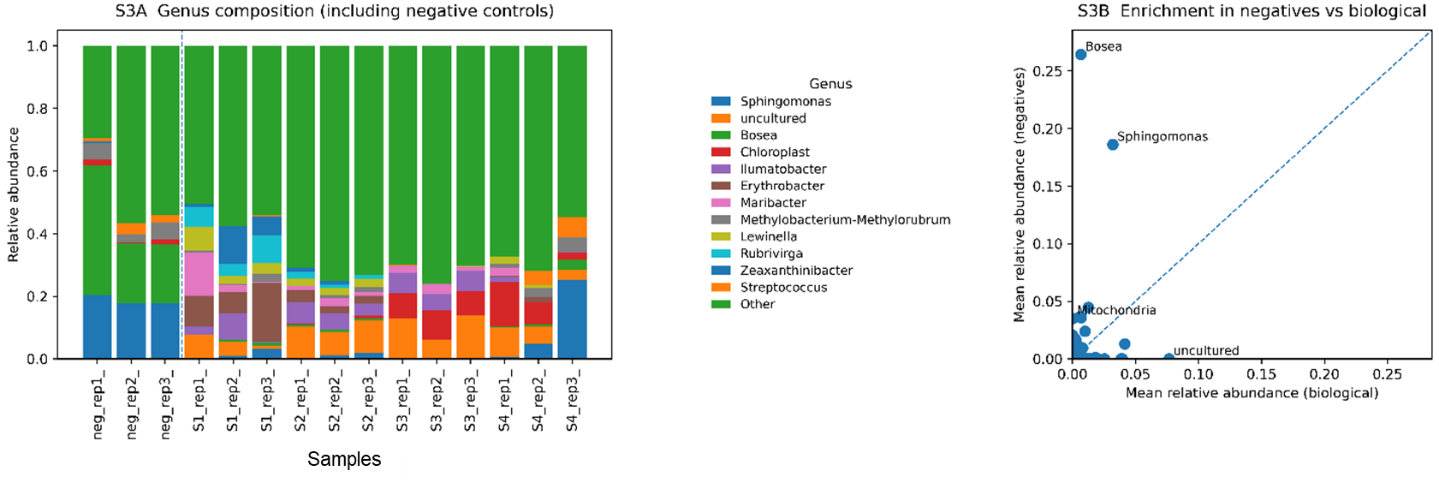

**Figure S3 | Negative-control–informed evaluation of background microbial signal.** (A) Genus-level relative abundance profiles across extraction negative controls and biological samples. Negative controls (left of dashed line) show enrichment of a restricted subset of low-biomass environmental taxa, whereas biological samples display broader compositional structure. Taxa are aggregated at the genus level, with low-abundance genera grouped as “Other.” (B) Mean genus abundance in negative controls versus biological samples. Each point represents a genus; the dashed diagonal denotes equal relative abundance between groups. Genera positioned above the diagonal are preferentially detected in negatives, consistent with background signal associated with reagents or laboratory environment, while most taxa remain low in both datasets, supporting minimal influence on the dominant biological community structure. S1: Flight_UV; S2: Flight_UVShield; S3:Ground_post; S4:Ground_pre; neg: No flight and UV.

### **1.4 Supplementary Methods S4. Evaluation of Filtering Impact on Ecological Structure**

To ensure that contaminant filtering did not artificially alter community structure, diversity patterns were compared before and after filtering. Principal coordinates analysis and diversity summaries demonstrated that major ecological gradients were preserved following filtering, supporting that the procedure reduced background signal without distorting biological relationships (Fig. S4).

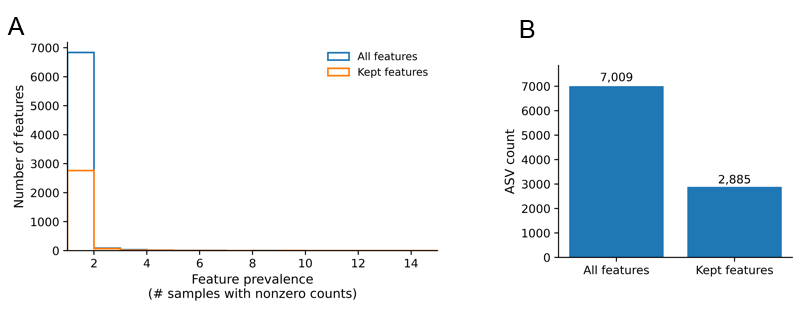

**Figure S4. Comparisons of diversity structure before and after filtering.** (A) Feature retention following prevalence-based filtering. The total number of amplicon sequence variants (ASVs) detected prior to filtering (“All features”, *n* = 7,009) and after application of the feature-selection criteria (“Kept features”, *n* = 2,885) are shown. Filtering reduced low-prevalence and potentially spurious features while preserving the majority of biologically relevant signal used in downstream diversity and compositional analyses. (B) Distribution of feature prevalence across samples before and after filtering. Histogram showing the number of samples in which each ASV was detected (non-zero counts) for the original feature table and the filtered dataset. Prior to filtering, most ASVs exhibited low prevalence across samples, consistent with rare or sparsely distributed taxa. The filtering procedure preferentially removed low-prevalence features, resulting in a modest rightward shift toward more consistently detected ASVs while retaining overall community structure.

#### **1.5 Supplementary Methods S5. Pairwise Multivariate Statistical Testing**

In addition to overall PERMANOVA tests reported in the main text, pairwise comparisons were performed to evaluate specific contrasts between experimental conditions, including UV exposure and ground-control timepoints. These analyses provide detailed statistical context for interpreting community separation observed in beta-diversity ordinations (Table S1).

**Table S1. Pairwise PERMANOVA results**

| **Comparison** | **Sample size** | **Pseudo-F** | **p-value** |
| --- | --- | --- | --- |
| Flight_UV vs. Flight_UVshield | 6 | 1.448542 | 0.110 |
| Flight_UV vs. Ground_post | 6 | 1.640377 | 0.097 |
| Flight_UVshield vs. Ground_post | 6 | 1.694701 | 0.096 |
| Groud_pre vs. Ground_post | 6 | 1.466228 | 0.104 |

#### **1.6 Supplementary Methods S6. Taxonomic Assignment Summary**

Homogeneity of multivariate dispersion was evaluated using PERMDISP based on Bray–Curtis distances with 999 permutations. Distances to group centroids were compared across conditions, and distributions were visualized as boxplots (Supplementary Fig. S5).

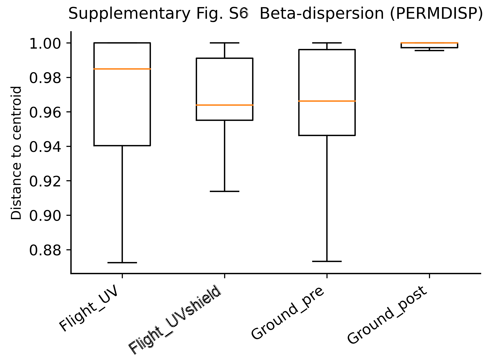

**Figure S5. Within-group beta-dispersion across experimental conditions.** Distributions of pairwise Bray–Curtis distances among samples within each condition (Flight_UV, Flight_UVshield, Ground_pre, Ground_post) are shown as boxplots. This analysis provides a complementary check that PERMANOVA results are not trivially driven by extreme within-group heterogeneity; formal PERMDISP statistics and pairwise tests are provided in the QIIME2 output.

To provide transparency regarding classification depth in low-biomass environmental samples, taxonomic assignment rates were summarized across hierarchical ranks. The proportion of sequences assigned to major taxonomic levels and the fraction remaining unclassified were evaluated to contextualize ecological interpretation and avoid overstatement of genus-level resolution (Fig. S6).

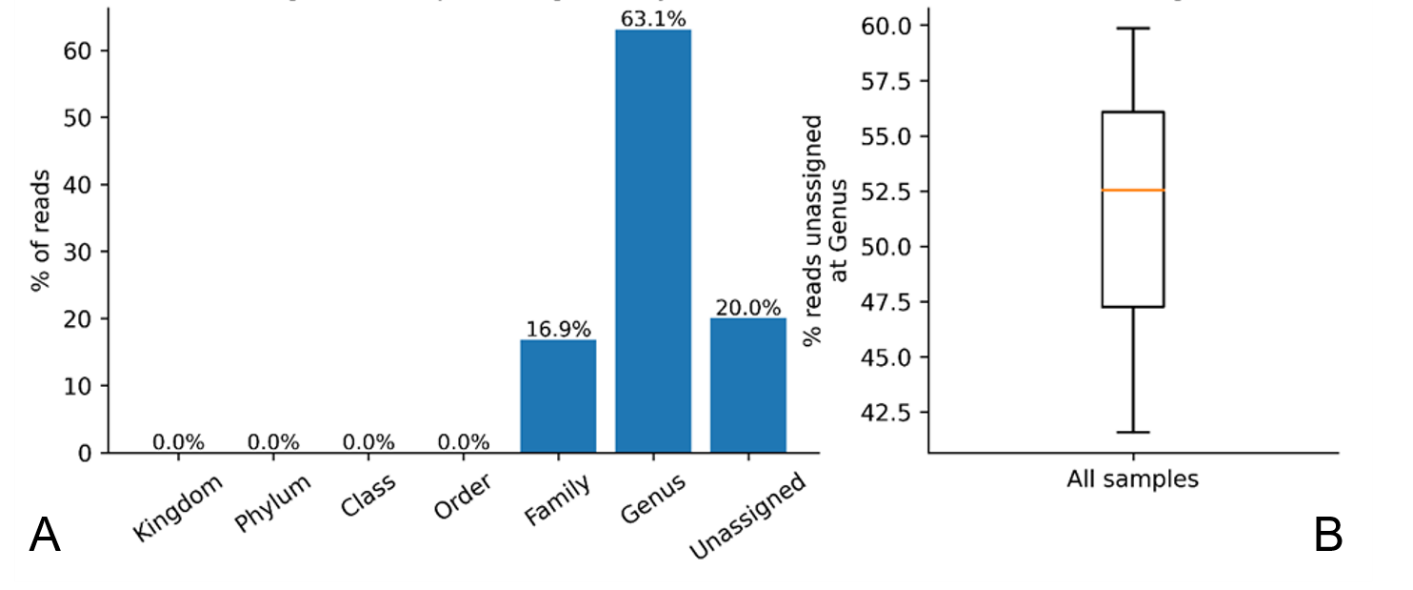

**Figure S6. Taxonomic assignment.** (A) Taxonomic assignment depth across hierarchical ranks. Bar plot showing the percentage of total sequencing reads assigned at each taxonomic rank based on SILVA reference classification, including Kingdom, Phylum, Class, Order, Family, Genus, and Unassigned categories. Assignment depth was determined using the deepest resolved taxonomic level for each amplicon sequence variant (ASV). This panel illustrates the overall classification resolution achieved in the dataset and provides transparency regarding the distribution of taxonomic confidence across ranks. (B). Fraction of genus-level unassigned reads per sample.
Boxplot summarizing the percentage of sequencing reads lacking genus-level assignment across all samples. The distribution highlights variability in classification depth among samples and reflects expected limitations of reference-database–based taxonomic annotation in complex environmental microbiomes. Outliers were not displayed to improve visualization clarity.

#### **1.7 Supplementary Methods S7. Differential abundance analysis (ANCOM)**

To identify microbial taxa exhibiting compositional differences among experimental conditions, differential abundance analysis was performed using the Analysis of Composition of Microbiomes (ANCOM) framework implemented in QIIME 2. ANCOM accounts for the compositional nature of amplicon sequencing data by evaluating pairwise log-ratio relationships among taxa rather than relying on absolute abundance estimates. Prior to analysis, a pseudocount was added to the feature table to accommodate zero values, and taxa were aggregated at the genus level to facilitate biological interpretation.

ANCOM was conducted using the experimental condition (Flight_UV, Flight_UVshield, Ground_pre, and Ground_post) as the grouping variable. The resulting W statistic represents the number of pairwise log-ratio tests for which the null hypothesis of equal abundance was rejected. Higher W values indicate stronger evidence that a taxon differs in relative abundance among experimental conditions. Differential abundance results were visualized using an ANCOM volcano plot and summarized by ranking taxa according to their W statistics.

The analysis identified multiple taxa with elevated W statistics, indicating that compositional shifts associated with near-space exposure were distributed across several phylogenetically distinct microbial lineages rather than being driven by a single dominant taxon. These findings complement the beta-diversity and PERMANOVA analyses by providing feature-level evidence of community restructuring. The twenty taxa with the highest ANCOM W statistics are presented in Supplementary Table S2.

**Table S2. Top 20 taxa ranked by ANCOM W statistic.** Taxa are ranked according to the ANCOM W statistic, representing the number of pairwise log-ratio comparisons supporting differential abundance among experimental conditions. Higher W values indicate stronger evidence for compositional differences across groups.

| **Rank** | **Genus** | **ANCOM W Statistic** |
| --- | --- | --- |
| 1 | d__Bacteria;p__Proteobacteria;c__Alphaproteobacteria;  o__Caulobacterales;f__Hyphomonadaceae;g__Hyphomonas | 333 |
| 2 | Unassigned | 325 |
| 3 | d__Bacteria;p__Proteobacteria;c__Alphaproteobacteria;  o__Sphingomonadales;f__Sphingomonadaceae;g__Sphingomicrobium | 320 |
| 4 | d__Bacteria;p__Desulfobacterota;c__Desulfobacteria;  o__Desulfobacterales;f__Desulfobacteraceae;g__Desulfoconvexum | 233 |
| 5 | d__Bacteria;p__Desulfobacterota;c__Desulfobulbia;o__Desulfobulbales;  f__Desulfocapsaceae;__ | 228 |
| 6 | d__Bacteria;p__Desulfobacterota;c__Desulfobulbia;o__Desulfobulbales;  f__Desulfocapsaceae;g__Desulforhopalus | 227 |
| 7 | d__Bacteria;p__Actinobacteriota;c__Actinobacteria;o__Propionibacteriales;  f__Nocardioidaceae;g__Nocardioides | 226 |
| 8 | d__Bacteria;p__Chloroflexi;c__Anaerolineae;o__SBR1031;f__A4b;g__A4b | 196 |
| 9 | d__Bacteria;p__Bacteroidota;c__Rhodothermia;o__Rhodothermales;  f__Rhodothermaceae;g__Rubrivirga | 186 |
| 10 | d__Bacteria;p__Bacteroidota;c__Bacteroidia;o__Bacteroidales;f__Prolixibacteraceae;  g__Draconibacterium | 181 |
| 11 | d__Bacteria;p__Desulfobacterota;c__Desulfobacteria;o__Desulfobacterales;  f__Desulfobacteraceae;g__Desulfobacter | 178 |
| 12 | d__Bacteria;p__Desulfobacterota;c__Desulfobulbia;o__Desulfobulbales;  f__Desulfocapsaceae;g__[Desulfobacterium]_catecholicum_group | 174 |
| 13 | d__Bacteria;p__Desulfobacterota;c__Desulfobulbia;o__Desulfobulbales;  f__Desulfocapsaceae;g__Desulfotalea | 173 |
| 14 | d__Bacteria;p__Proteobacteria;c__Alphaproteobacteria;o__Rhizobiales;__;__ | 167 |
| 15 | d__Bacteria;p__Bacteroidota;c__Bacteroidia;o__Cytophagales;f__Cyclobacteriaceae;  g__Marinoscillum | 166 |
| 16 | d__Bacteria;p__Proteobacteria;c__Alphaproteobacteria;o__Rhodobacterales;  f__Rhodobacteraceae;g__Rhodobacteraceae | 164 |
| 17 | d__Bacteria;p__Bacteroidota;c__Bacteroidia;o__Flavobacteriales;f__Flavobacteriaceae;  g__Algibacter | 162 |
| 18 | d__Bacteria;p__Proteobacteria;c__Alphaproteobacteria;o__Sneathiellales;  f__Sneathiellaceae;g__Sneathiella | 161 |
| 19 | d__Bacteria;p__Desulfobacterota;c__Desulfobacteria;o__Desulfobacterales;  f__Desulfobacteraceae;g__Desulfobacterium | 158 |
